# Glutaminase contributes to MYC-induced cell-autonomous autophagy and to Ras^V12^-dependent non-autonomous autophagy in the Drosophila wing disc epithelium

**DOI:** 10.64898/2026.08.31.748041

**Authors:** Francesca Destefanis, Lorenzo Conci, Shivani Bajaj, Valeria Manara, Paola Bellosta

## Abstract

MYC-driven metabolic reprogramming supports rapid cell growth but also creates metabolic demands that require adaptive mechanisms to maintain cellular homeostasis. Here, combining clonal analysis in *Drosophila* wing imaginal discs with studies in Schneider S2 cells, we identify glutamine metabolism as a component of Myc-induced autophagy. Myc increased the expression of genes involved in glutamine utilization, including glutaminase (GLS), and enhanced ammonia production, a metabolic by-product of glutaminolysis. Genetic depletion of GLS in clones suppressed the accumulation of Myc-induced Atg8a-positive structures and reduced autophagic flux, demonstrating that glutaminase contributes to the autophagic response elicited by Myc. Exogenous NH₄Cl was sufficient to induce Atg8a-positive structures and partially restored their accumulation following GLS depletion, supporting ammonia as a downstream contributor to this response. Mechanistically, Myc-induced autophagy in clones required the core autophagy factor Atg5 but was not suppressed by activation of Rheb/TOR signaling or Atg1 depletion, indicating reduced dependence on canonical TOR-Atg1 regulation. We further found that Myc activity is required for Ras^V12^-driven epithelial overgrowth and that Ras^V12^ cells induce a pronounced non-cell-autonomous accumulation of Atg8a-positive structures in wild-type cells surrounding Ras^V12^ clones. Depletion of either Myc or Gls in Ras^V12^ cells strongly reduced this neighboring autophagic response. Together, our findings identify Gls-dependent glutamine metabolism as a previously unrecognized component of Myc-induced autophagy and extend this relationship to Ras-transformed epithelia, linking the metabolic state of transformed cells to autophagy in the surrounding tissue.

**Graphical abstract.:** 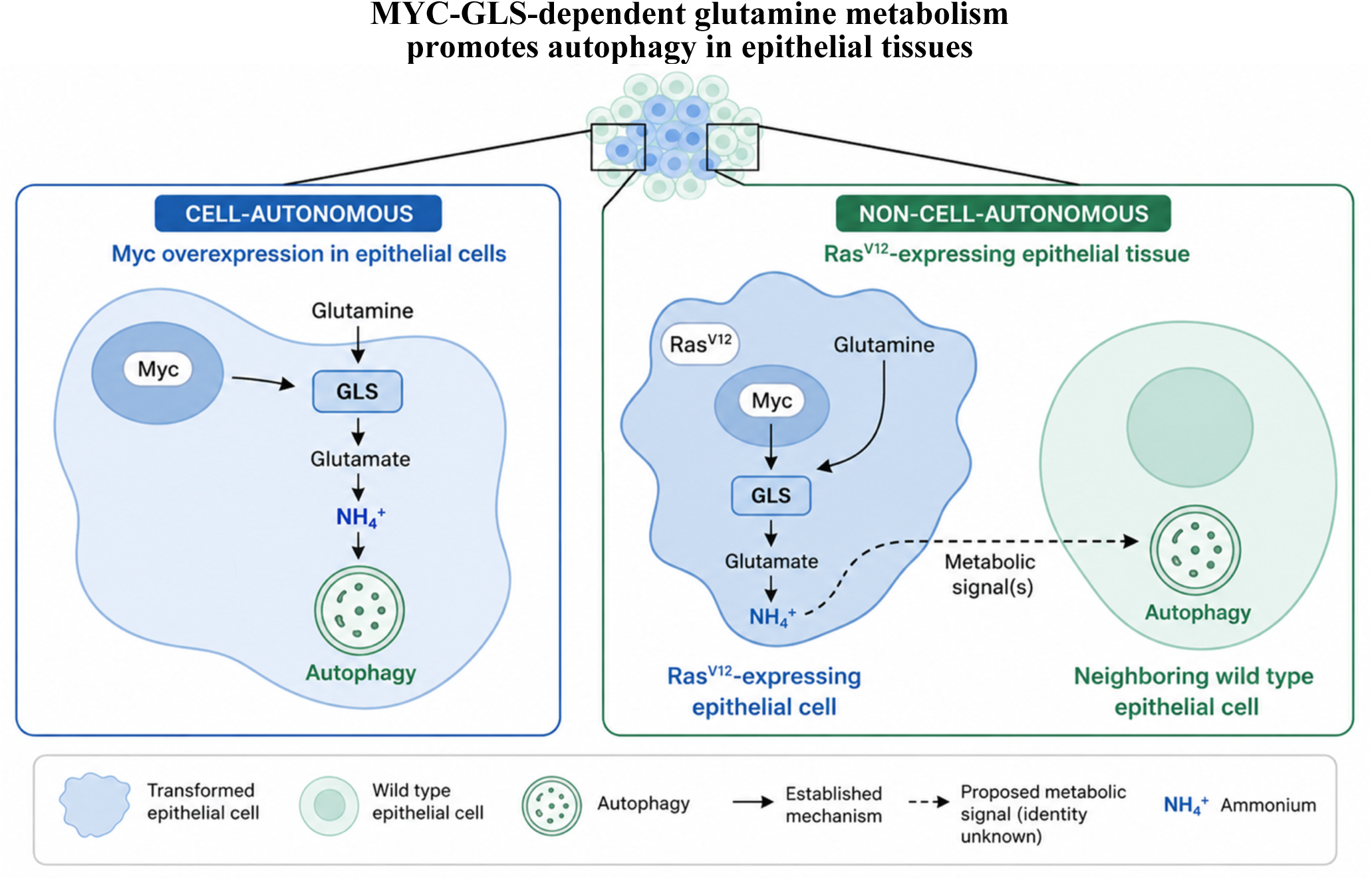

Myc increases glutaminase (Gls)-dependent glutamine catabolism, promoting ammonia production and Atg5-dependent autophagy in Drosophila epithelial cells. In Ras^V12-^transformed epithelia, Myc and Gls are also required for the induction of autophagy in neighboring wild-type cells, suggesting that metabolic signals generated by transformed cells can elicit a non-cell-autonomous autophagic response. Solid arrows indicate experimentally supported relationships, whereas the dashed arrow denotes a proposed metabolic signal whose identity remains to be established.

## 1. Introduction

MYC is an evolutionarily conserved transcription factor that coordinates cell growth, proliferation, and metabolism [1, 2]. Among its metabolic functions, MYC promotes glutamine uptake and utilization by regulating amino acid transporters and glutaminase (GLS), thereby supporting TCA-cycle anaplerosis and anabolic metabolism [3, 4]. This dependence on glutamine metabolism is particularly relevant in cancer, where MYC-driven glutaminolysis supports the metabolic requirements of tumor cell growth and survival and can create therapeutically exploitable metabolic vulnerabilities [5, 6]. MYC-dependent regulation of genes involved in glutamine metabolism is also observed in *Drosophila*. We previously showed that Myc promotes the expression of key components of glutamine metabolism and amino acid transport in the metabolic fat body, and in the *Drosophila* Schneider-S2 cell-based model of Myc-driven cell competition [7, 8]. Among these metabolic targets, glutaminase is of particular interest because its conversion of glutamine to glutamate generates ammonia, a metabolite with established autophagy-inducing activity. Importantly, ammonia has been shown to induce autophagy through a TOR/ULK1-independent mechanism in mammalian cells [9–12], suggesting that glutamine catabolism may directly couple metabolic rewiring to non-canonical autophagic response.

Autophagy represents an important adaptive response to the metabolic and proteotoxic demands associated with elevated MYC activity. In mammals, the relationship between MYC and autophagy is highly context dependent, with MYC reported to either suppress the autophagy–lysosomal program or promote autophagy as part of the metabolic adaptations that sustain MYC-driven cells [13]. MYC-dependent metabolic rewiring has been linked to glutamine catabolism in prostate cancer, where MYC-driven GLS activity promotes ATG5-dependent autophagy, helping cancer cells control oxidative stress, maintain stem-like properties, and resist treatment [14]. More broadly, MYC-induced autophagy acts mainly through stress-responsive pathways that can operate independently of the canonical TOR and the Atg1/ULK1 pathway [15]. Consistent with this adaptive role, MYC-induced autophagy was already linked to ER stress and the PERK-eIF2α-ATF4-dependent pathway rather than to mTOR signaling [16]. Similarly, in *Drosophila,* Myc-driven growth induces UPR-associated autophagy together with Ref2(P)/p62/SQSTM1, dependent activation of the antioxidant transcription factor Nrf2 [17]. We further showed that Myc-induced autophagy is coupled to metabolic remodeling, with lipid metabolism representing an important component of the autophagic response that supports Myc-driven tissue growth [18].

Beyond supporting cell-autonomous adaptation, MYC-dependent metabolic programs can also influence how cells interact with their surrounding tissue. Myc is a key regulator of physiological cell competition, an evolutionarily conserved process first characterized in *Drosophila* [19, 20] and subsequently demonstrated in mammalian tissues, where it contributes to development and tissue homeostasis [21–23]. Cell competition also influences the fate of transformed cells in both *Drosophila* and mammalian epithelia [24]. In mammalian epithelial models, for example, Ras^V12^-expressing cells can be recognized and eliminated by surrounding wild-type cells through apical extrusion [25, 26], whereas in *Drosophila,* oncogenic and metabolic alterations can modify competitive interactions and promote tumor growth [27].

Previous studies demonstrated that Myc levels increase in Ras^V12^-driven overgrowth and contribute to the competitive behavior of transformed cells [28]. Given the central role of Myc in coordinating metabolism and cellular fitness [29], it remains an important unresolved question whether Myc-dependent glutamine metabolism contributes to Ras-driven tumor growth and to autophagic responses in neighboring cells. In *Drosophila*, Ras^V12^-driven tumor models recapitulate several hallmarks of epithelial tumorigenesis, including tissue overgrowth and extensive interactions with surrounding wild-type cells, which, in combination with the loss of polarity tumor suppressors, such as *dlg*, *scrib*, or *lgl*, drives malignant tumor growth and invasion [30–32]. In particular, in a Drosophila tumor model based on cooperation between RasV12 and loss of scribble (scrib), non-cell-autonomous autophagy is induced in the surrounding microenvironment and supports tumor growth, in part by increasing nutrient availability to transformed cells [33, 34]. However, how the metabolic state of transformed cells contributes to the induction of this non-cell-autonomous autophagic response remains less understood.

Here, we identify Gls-dependent glutamine metabolism as a component of Myc-induced autophagy in *Drosophila* epithelial cells. Myc increased Gls expression and ammonia production, whereas Gls depletion impaired Myc-induced autophagic flux. Exogenous ammonia promoted the accumulation of Atg8a-positive vesicles and partially restored this response following Gls depletion. Genetic analysis further showed that Myc-induced autophagy requires Atg5 but displays reduced dependence on canonical Rheb/TOR-Atg1 regulation. Finally, in Ras^V12^-transformed epithelia, Myc and Gls activity within transformed cells was required to induce autophagy in neighboring wild-type tissue. Together, these findings connect MYC-dependent glutamine metabolism to both cell-autonomous and non-cell-autonomous regulation of autophagy.

## 2. Experimental Procedures

### 2.1 Drosophila stocks and husbandry

*Drosophila* stocks were maintained under standard culture conditions on conventional cornmeal-molasses-based medium at 25°C unless otherwise indicated. Genetic crosses were performed using standard procedures. The GAL4/UAS system was used for transgene expression, and FLP/FRT-based approaches were used to generate genetically defined mosaic tissues.

The following transgenes, alleles, and reporters were used in this study: the ey>d*myc^wt^* and hypomorphic ey>d*myc^P0^* lines [35]; Actin>CD2>Gal4, UAS-GFP a gift from Bruce Edgar, University of Utah Salt Lake City, UT; Tubulin>CD2>Gal4, UAS-GFP a gift from Eugenia Piddini, University of Bristol, Bristol, UK; MS1096>Gal4 (BDSC#8860); UAS-HA-Myc [35]; UAS-Gls-RNAi (VDRC# 108317, # 7192); UAS-Myc-RNAi (VDRC#2747 BDSC#25783); UAS-Atg1-RNAi and UAS-Atg5-RNAi a gift from Thomas Neufeld, University of Minnesota, Minneapolis, MN; UAS-Ras^V12^, UAS-dlg-RNAi a gift from Daniela Grifoni, University of L’Aquila, IT [36]; UAS-Rheb^AV4^ gift from Hugo Stocker, ETH, Zurich CH [37]; UAS-Luciferase-RNAi (BDSC#31603); UAS-mCherry-Atg8a-GFP reporter line (BDSC#37749); 3x mCherry-GFP-Atg8a [38] and Tubulin-Ref(2)P-GFP [39] a gift from Juhász Gabor. Two independent RNAi lines targeting either *Gls* or *myc* were used throughout the study, and their knockdown efficiency was validated by qRT–PCR (Supplementary Figure 1). UAS-Atg1 and Atg5 RNAi lines were validated in [40].

### 2.2 Generation of FLP-out clones in wing imaginal discs

FLP-out recombination was induced at 48 h after egg laying (AEL), following a 2-4 h egg-collection period, by heat shock at 37°C in a water bath. Larvae carrying hs-FLP^122^; Actin>CD2>Gal4, UAS-GFP [28] or hs-FLP^122^; Tubulin>CD2>Gal4, UAS-GFP were heat-shocked for 20-30 minutes and subsequently maintained under standard conditions until dissection at the third-instar stage. In all the experiments, the hs-FLP^122^ allele on the X chromosome was used [41],

### 2.3 Wing imaginal disc dissection and immunofluorescence

Wing imaginal discs from third-instar larvae were dissected in PBS 1X and fixed in 4% PFA (EMS #15710) for 30 minutes at room temperature on a rotator. Tissues were washed in PBS 1X, permeabilized with PBS/0.3% Triton, washed again in PBS/0.01% Tween, and blocked with PBS/3% BSA before incubation with the indicated primary and secondary antibodies, as required. Nuclei were visualized with Hoechst 33352, where indicated. Samples were mounted in Vectashield (H-1900 Vector Laboratories) and imaged using an SP8 Leica confocal microscope. Equivalent acquisition settings were used for samples included in the same quantitative comparison.

### 2.4 Quantification of Atg8a-positive structures in FLP-out clones

For experiments examining Myc- and Gls-dependent autophagy and to determine the genetic requirements for Myc-induced autophagy, GFP-marked FLP-out clones were generated to express Myc alone or in combination with GLS-RNAi and with Rheb^AV4^, Atg1-RNAi, or Atg5-RNAi. Wing imaginal discs containing GFP-labeled FLP-out clones expressing Atg8a-positive puncta within GFP-positive clones were quantified using the PECAN image-analysis pipeline [42]. GFP-positive clones were identified and used as regions of interest (ROIs). The plugin was used to detect Atg8a-positive puncta using identical analysis parameters for all genotypes. The total area occupied by Atg8a-positive puncta was calculated and normalized to the corresponding GFP-positive clone area. Results are presented as the percentage of clone area occupied by Atg8a-positive structures. Multiple clones from at least three independent imaginal discs were analyzed for each genotype. Statistical significance was assessed using one-way ANOVA followed by Tukey’s multiple comparisons test. For each biological experiment, 3–5 wing imaginal discs were analyzed per genotype, with approximately four clones quantified per disc. Data were obtained from two or three independent biological experiments, each containing at least five to six wing imaginal discs per genotype. Confocal images were acquired using identical acquisition settings for all samples within an experiment.

A similar protocol was used to induce FLP-out clones, which were used to quantify Atg8a puncta following NH_4_Cl treatment in the rescue experiments shown in Figure 2. Image analysis was performed from confocal z-stacks, using Fiji/ImageJ2 on GFP-positive clones. A common fluorescence threshold was applied to all images to identify Atg8a-positive structures. The integrated density of thresholded Atg8a-positive puncta was measured in three fixed-size regions of interest (ROIs) per clone. Identical thresholding parameters and ROI dimensions were applied to all samples within an experiment. Measurements obtained from multiple clones were averaged to generate a value for each biological replicate. Two independent biological experiments were analyzed. Data shown represent the mean values from independent experiments. Statistical significance was determined using one-way ANOVA followed by Tukey’s multiple-comparison testing.

**Figure 1.**
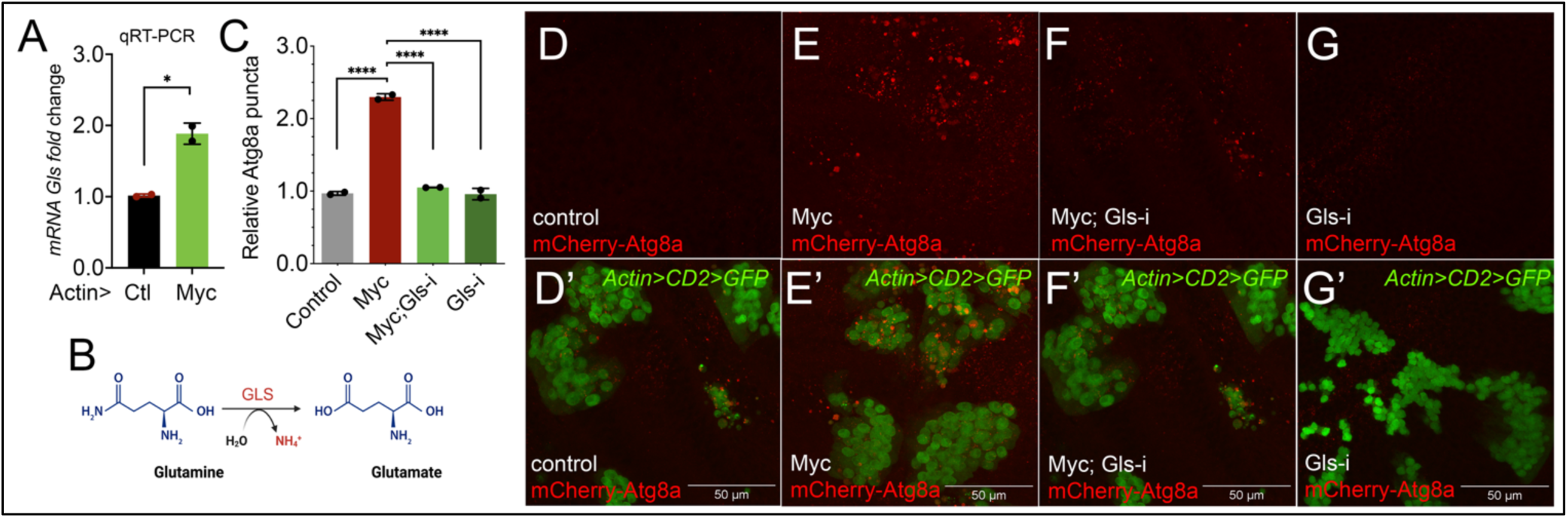
Myc promotes Gls expression and Gls-dependent accumulation of Atg8a-positive vesicles. (A) qRT-PCR analysis of *glutaminase (Gls) mRNA* in wing imaginal discs ubiquitously overexpressing Myc under the actin promoter. *Gls* expression was normalized to *Actin5C,* and fold change was calculated relative to the control. Data are from two independent biological experiments, each using 30-50 wing imaginal discs. Statistical significance was determined using Student’s t-test. \**P* < 0.05. (B) Schematic representation of the GLS-catalyzed conversion of glutamine to glutamate, resulting in the production of ammonium (NH₄⁺). (C) Quantification of Atg8a-positive puncta in FLP-out clones of the indicated genotypes. Atg8a-positive puncta were quantified using the PECAN image-analysis pipeline and normalized to the mean value of control clones, using 3–5 wing imaginal discs per genotype, with an average of four clones quantified per disc in each biological experiment. Data are from two independent biological experiments. Bars indicate mean ± SEM. Statistical significance was assessed by one-way ANOVA followed by Tukey’s multiple-comparisons test, \*\*\*\**P* < 0.0001. (D–G′) Representative images of FLP-out clones expressing GFP alone (Control; D, D′), Myc (E, E′), Myc together with Gls-RNAi (Myc; Gls-i; F, F′), or Gls-RNAi alone (Gls-i; G, G′), in wing imaginal discs carrying the autophagy reporter mCherry-Atg8a in red and GFP-positive FLP-out clones in green. (D–G) mCherry-Atg8a channel; (D′–G′) merged mCherry-Atg8a and GFP images. Scale bars, 50 μm.

**Figure 2.**
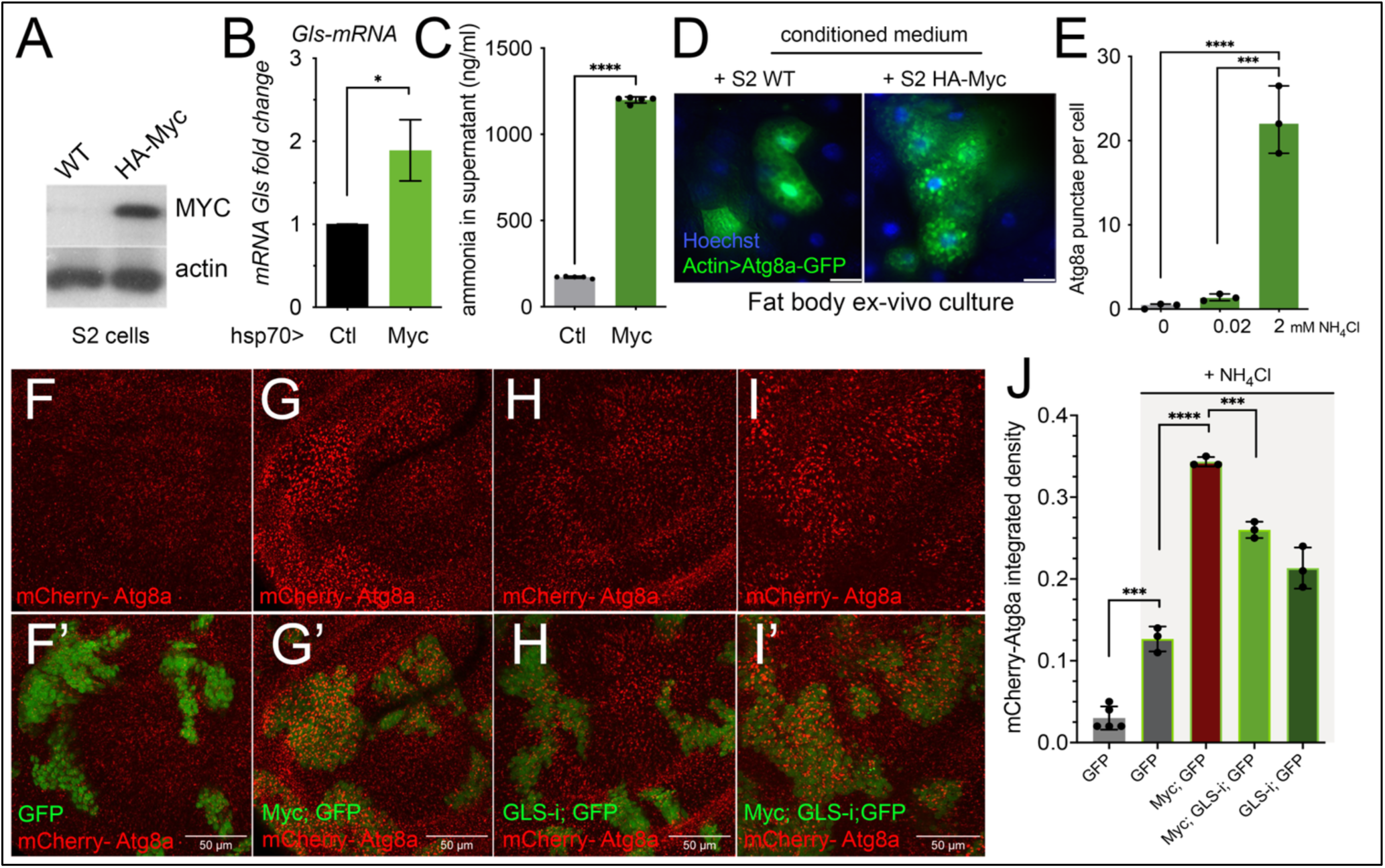
Exogenous ammonia promotes Atg8a accumulation and partially rescues Gls depletion. (A) Representative immunoblot confirming HA-Myc expression in S2 cells. HA-Myc was detected using an anti-HA antibody, and actin was used as a loading control. (B) RT-qPCR analysis of Gls mRNA in control and Myc-expressing S2 cells. Myc expression was induced from the hsp70 promoter, and transcript levels were normalized to Actin5C and expressed as fold change relative to control. Statistical significance was determined using Student’s t-test; *\*P* < 0.05. (C) Ammonia concentration in the culture medium of control (WT) and HA-Myc-expressing S2 cells, measured using a colorimetric ammonia assay. Data are shown as mean ± SEM. Statistical significance was determined using an unpaired two-tailed Student’s t-test; \*\*\*\**P* < 0.0001. (D) Representative images of larval fat bodies carrying Atg8a-GFP-expressing FLP-out clones and cultured ex vivo for 4 h in conditioned medium collected from control (WT) or HA-Myc-expressing S2 cells. Nuclei are labeled with Hoechst 33352 (blue), and Atg8a-GFP-positive vesicles are shown in green. (E) Quantification of Atg8a-positive puncta per cell in ex vivo cultures of fat bodies carrying FLP-out clones from control animals following treatment for 4 h with the indicated concentrations of NH₄Cl. Statistical significance was determined using one-way ANOVA followed by multiple-comparison testing; \*\*\**P* < 0.001; \*\*\*\**P* < 0.0001. (F–I) Representative confocal images of GFP-marked FLP-out clones expressing the indicated transgenes in wing imaginal discs following ex vivo incubation for 4 h with 2 mM NH₄Cl in Schneider medium. mCherry-Atg8a is shown in red and GFP-marked clones in green. (J) Quantification of mCherry-Atg8a-positive puncta in GFP-marked clones of the indicated genotypes. NH₄Cl treatment increased Atg8a-positive puncta in control clones, and this accumulation was further increased in Myc-expressing clones. Gls depletion reduced Atg8a accumulation in Myc-expressing clones, while NH₄Cl partially restored Atg8a accumulation in Myc; Gls-RNAi clones. Quantification was performed by measuring the integrated density of thresholded mCherry-Atg8a-positive puncta in three fixed-size ROIs per clone across three independent experiments using Fiji/ImageJ. Identical thresholds and ROI dimensions were applied to all samples. Measurements from multiple clones were averaged for each biological replicate. Data are shown as mean ± SEM. Statistical significance was determined using one-way ANOVA followed by multiple-comparison testing; \*\*\**P* < 0.001; \*\*\*\**P* < 0.0001. Scale bars, 50 μm.

### 2.5 RNA extraction and quantitative RT-PCR

For in vivo analysis of Gls expression, RNA was extracted from wing imaginal discs ubiquitously expressing Myc under the control of an Actin-Gal4 promoter and from the corresponding controls. Each biological replicate contained approximately 30–50 wing imaginal discs.

For experiments in Schneider S2 cells, RNA was isolated from a small flask of S2-wt and from S2-hsp70-HA-Myc-expressing cells, in which Myc overexpression was achieved by heat shock (see next). Total RNA was isolated using the Qiagen-RNeasy mini kit, and cDNA was synthesized from 1µg of total RNA using Superscript IV-VILO (#11754050 Invitrogen). Quantitative PCR was performed using PowerTrack SYBR Green Master Mix (Thermo Fisher Scientific). Transcript abundance was normalized to Actin5C, and expression is reported as fold change relative to the corresponding control. Primer sequences are provided in Supplementary Table 1.

### 2.6 *Drosophila* Schneider S2 cell culture and induction of Myc expression

*Drosophila* Schneider S2 cells were grown at 25°C in flasks using Schneider medium (Gibco) supplemented with 10% heat-inactivated fetal calf serum (FCS) and 100 IU of penicillin-streptomycin (Gibco). S2-hsp70-HA-Myc cell line used in Figure 2, was transfected with pCasper-hsp-HA-Myc using the pcP4 plasmids carrying the heat-shock *hsp70*-inducible promoter, using Cellfectin reagent (Invitrogen) as described in [35]. For inducible Myc expression, S2-HA-Myc were subjected to a heat-shock by incubation for 30 minutes at 37°C, and protein expression was analyzed two hours after heat-shock by western blot. Control cells were treated in parallel. The *Drosophila* Schneider S2-mt-HA-Myc cell line used in Figure 3 was generated by transfection with the HpRmHa-3-HA-MYC plasmid [43], in which HA-tagged Myc expression is driven by the metallothionein promoter. Myc expression was induced by adding CuSO₄ to the culture medium at a final concentration of 0.7 mM for 4 h prior to RNA extraction and Western blot analysis.

**Figure 3.**
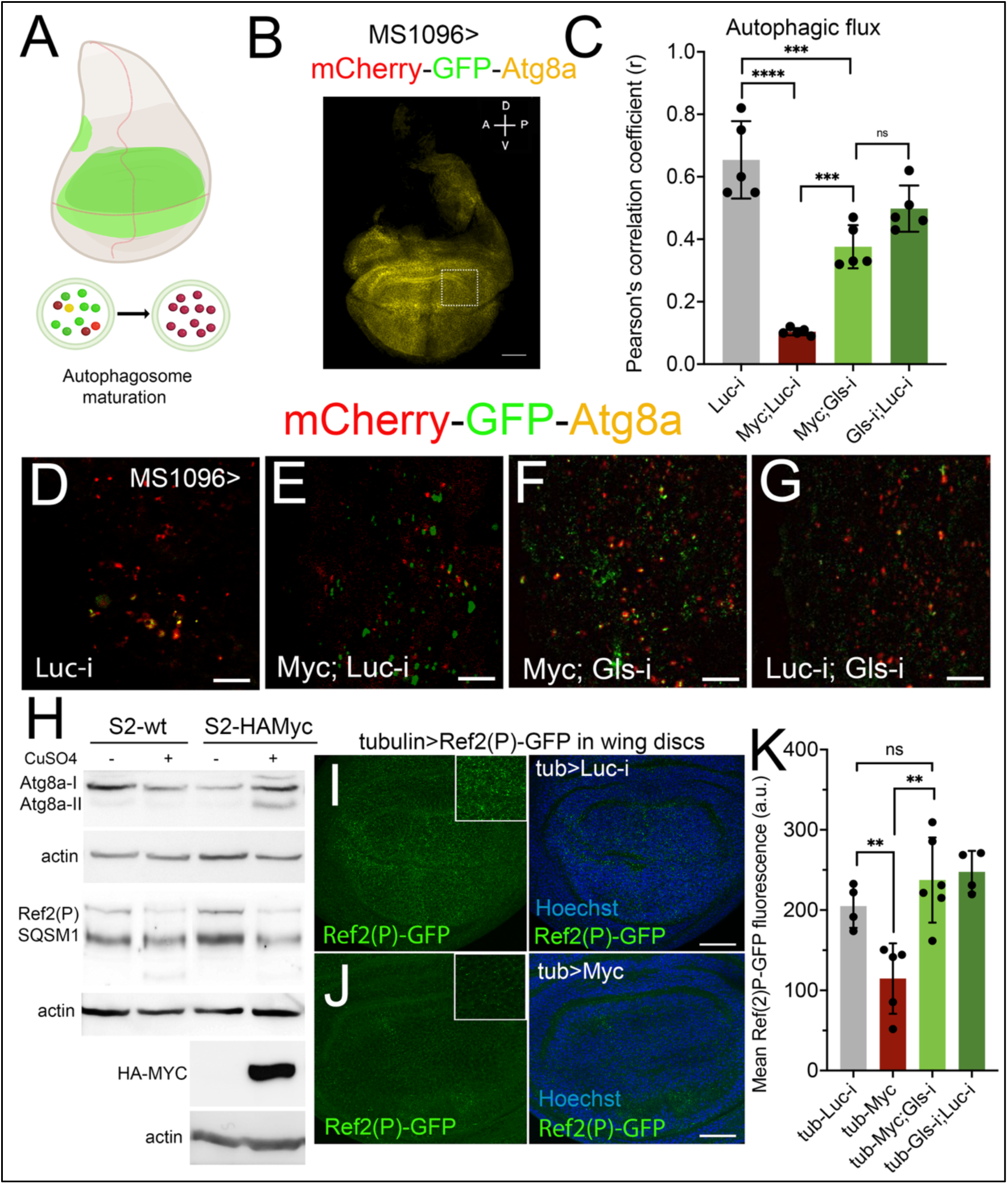
Myc promotes autophagic flux in *Drosophila* wing discs in a glutaminase-dependent manner. (A) Schematic representation of the expression domain of the MS1096-Gal4 driver in the wing pouch (green) and of autophagosome maturation monitored using the tandem mCherry-GFP-Atg8a reporter. Newly formed autophagosomes are positive for both GFP and mCherry (yellow), whereas following fusion with lysosomes, GFP fluorescence is quenched in the acidic environment while mCherry remains stable, resulting in red-only autolysosomes indicative of productive autophagic flux. (B) Representative confocal image of an MS1096>mCherry-GFP-Atg8a wing imaginal disc. Scale bar, 50 μm. The white-boxed region indicates the area used for high-magnification analysis, as shown in (D–G) D-dorsal, V-ventral, and A-antero, P-posterior axis. (C) Quantification of autophagic flux by Pearson’s correlation coefficient (r) between GFP and mCherry fluorescence. Myc overexpression significantly reduced GFP/mCherry colocalization compared with controls, indicating increased autophagic flux, whereas glutaminase (Gls) reduction partially restored colocalization, demonstrating that Myc-induced autophagic flux requires glutamine metabolism. Data are shown as mean ± SEM; each dot represents one wing disc *\*\*\*P* < 0.001, *\*\*\*\*P* < 0.0001. (D–G) Representative high-magnification images of the regions analyzed for Pearson’s correlation. Scale bars, 10 μm. (D) Control (Luc-i), (E) Myc overexpression (Myc; Luc-i), (F) Myc overexpression combined with Gls-RNAi (Myc; Gls-i), and (G) Gls RNAi combined with control transgene Luc-RNAi (Gls-i; Luc-i). Red puncta correspond to acidified autolysosomes, whereas yellow puncta represent GFP- and mCherry-positive autophagosomes. (H) Immunoblot analysis of endogenous Atg8a and Ref(2)P/SQSTM1 in control (S2-wt) and inducible HA-Myc-expressing S2 cells following CuSO₄ treatment. Myc expression increased the abundance of lipidated Atg8a-II and reduced Ref(2)P protein levels, consistent with enhanced autophagic activity and cargo turnover. Densitometric quantification of Atg8a lipidation is shown below the corresponding lanes as the Atg8a-II/Atg8a-I ratio, normalized to untreated WT cells = 1, and Ref(2)P levels, normalized to Actin in untreated WT = 1, are indicated. Actin served as the loading control. HA-Myc expression was confirmed by immunoblotting. (I–J) Representative images of Tubulin>Ref(2)P-GFP expression in wing discs co-expressing Luc-i (I) or Myc (J). Ref(2)P-GFP fluorescence is reduced upon Myc overexpression, consistent with increased turnover of the selective autophagy cargo receptor. Insets show enlarged regions of the pouch. Scale bars, 50 μm. (K) Quantification of mean Ref(2)P-GFP fluorescence in wing discs. Myc expression significantly reduced Ref(2)P-GFP levels compared with controls, whereas simultaneous Gls reduction restores Ref(2)P-GFP fluorescence near control levels, indicating that Gls is required for the Myc-dependent reduction of Ref(2)P. These findings are consistent with enhanced autophagic turnover and are supported by the tandem mCherry-GFP-Atg8a autophagic flux reporter (panel C). Bars represent mean ± SEM; each dot represents one wing disc. Statistical significance was determined using an unpaired two-tailed Student’s t-test *\*\*P* < 0.01.

### 2.7 Measurement of ammonia production

Ammonia production was measured in control and Myc-expressing S2 cells using a colorimetric ammonia assay kit (Bio Vision, #K370). Following induction of Myc expression in 80% confluent flasks, the supernatant was collected on day 2 of induction. Medium was cleared of cells and debris by centrifugation and used immediately. Ammonia concentrations were determined according to the manufacturer’s instructions. Absorbance was measured at 570 nm using a plate reader, and ammonia concentrations were calculated from a standard curve. Values were normalized to S2-WT cells treated in parallel.

### 2.8 Preparation of fat bodies in ex vivo conditions

For ex vivo experiments, larval fat bodies containing GFP-marked FLP-out clones and expressing Atg8a-GFP autophagy reporter were dissected in complete Schneider’s medium supplemented with 10% FBS and incubated for 4 h with conditioned medium obtained from control or HA-Myc-expressing S2 cells. Atg8a-GFP autophagic vesicles were visualized using a Zeiss Axio Imager M2 fluorescence microscope (Carl Zeiss Microscopy).

### 2.9 Ex vivo NH_4_Cl treatment

Third instar larval fat bodies were dissected in complete Schneider medium. For dose-response experiments, larval fat bodies were incubated for 4 h with increasing concentrations of NH4Cl (0 mM, 0.02 mM, and 2mM). For wing imaginal disc experiments, dissected discs carrying GFP-marked FLP-out clones and the mCherry-Atg8a reporter were incubated for 4 h in complete Schneider medium in the absence or presence of 2 mM NH4Cl. Following incubation, tissues were processed and imaged under identical conditions.

### 2.10 Tandem mCherry-GFP-Atg8a autophagic flux analysis

Autophagic flux in wing imaginal discs was assessed using the tandem UAS-mCherry-GFP-Atg8a reporter expressed in the wing pouch under the control of MS1096-Gal4 promoter.

GFP and mCherry fluorescence intensities were measured within the MS1096-Gal4 expression domain using Fiji/ImageJ2. Colocalization was analyzed using intensity correlation analysis, and the Pearson correlation coefficient between GFP and mCherry signals was calculated. A reduction in GFP/mCherry correlation was interpreted as increased delivery of Atg8a-positive structures to acidic compartments and, therefore, increased autophagic flux. Wing imaginal discs from approximately 5-10 animals per genotype were analyzed.

### 2.11 Ref(2)P-GFP analysis

The Tubulin>Ref(2)P-GFP reporter was used to analyze the autophagic cargo turnover in wing imaginal discs. This line expresses the Ref(2P)-GFP construct from Neufeld’s lab [39] cloned under the tubulin promoter from Gabor’s laboratory. Ref(2)P-GFP fluorescence was quantified from confocal stacks within the wing disc pouch using Fiji/ImageJ2. Identical acquisition settings, thresholding parameters, and ROI dimensions were applied to all samples within each experiment. Three measurements from multiple ROIs within each wing disc were averaged to obtain a single value per biological replicate. Two independent biological experiments were performed. Statistical significance was determined by one-way ANOVA followed by Tukey’s multiple-comparison test. Reduced Ref(2)P-GFP fluorescence was interpreted as increased autophagic turnover.

### 2.12 Immunoblot analysis

Protein extracts from control and inducible S2-HA-Myc cells were prepared in lysing buffer (50 mM Hepes pH 7.4, 150 mM NaCl, 1.5% Triton, 1mM EDTA), with phosphatase and protease inhibitors (Roche). Protein concentrations were quantified using Bradford reagent. Equal amounts were resolved on SDS-PAGE gels and transferred to nitrocellulose membranes. After blocking with 5% non-fat milk in 1X TBS-Tween 0.01% (TBST), membranes were incubated overnight at 4°C with the primary antibody, rat anti-HA (for HA-Myc), at 1:1000 (MoAb 3F10 Roche #11867423001), mouse anti-actin at 1:400 (DSHB JLA20), rabbit anti-Ref(2)P at 1:500 (ABCAM#178440), and rabbit anti-ATG8a at 1:800 (MERCK ABC974). After washing in TBST, membranes were incubated with the specific HRP-conjugated secondary antibodies (Cell Signaling Technology, diluted 1:1000) for 2 hours, then washed with TBST, and enhanced chemiluminescence (ECL, Amersham, Merck) was used for detection. Atg8a-I and lipidated Atg8a-II species were quantified to determine the Atg8a-II/Atg8a-I ratio. Ref(2)P levels were normalized to the corresponding loading control.

### 2.13 Ras^V12^-driven tumor models and quantification of Ras^V12^-induced tissue overgrowth in a *myc* **mutant background.**

Ras-driven tissue overgrowth was examined using an eyeless-FLP-based recombination system to generate mosaic tissues expressing Ras^V12^ alone or Ras^V12^ together with dlg-RNAi in the eye-antennal under the *eyeless* compartment [35]. The requirement for Myc activity was examined by generating these tissues either in animals carrying wild-type Myc activity (*ey>dmyc^WT^*) or in animals hemizygous for the hypomorphic *dmyc^P0^* allele (*ey>dmyc^P0^*).

Third-instar larval eye-antennal imaginal discs were dissected and processed for fluorescence microscopy. GFP was used to visualize genetically manipulated tissue. Eye-antennal imaginal disc area was measured from confocal images using Fiji/ImageJ2. Values were normalized to the mean area of the corresponding *ey>dmyc^WT^* control discs. Each data point represents an individual animal. Where indicated, differentiated photoreceptor cells were detected using anti-Elav as 1:100 (DSHB #9F8A9), and nuclei were labeled with Hoechst 33352.

### 2.14 Analysis of non-cell-autonomous autophagy surrounding Ras^V12^ clones

To investigate autophagic responses associated with Ras-driven cell competition, in which Ras^V12^-transformed cells can be outcompeted and eliminated by surrounding wild-type cells, GFP-marked Ras^V12^ FLP-out clones were generated and analyzed in wing imaginal discs carrying the ubiquitously expressed mCherry-Atg8a reporter using a Gal80^ts^-controlled clonal system. Animals were left laying eggs for 3-4 hours in standard fly food at 25°C, then heat shock was performed at 37°C in a water bath on F1 larvae at 48 or 72 hours AEL for 12 minutes (for hs-flp122; tubulin>Gal80ts, y+>Gal4; UAS-GFP/SM5-TM6b) or 10 minutes (for w; tubulin-Gal80ts; actin>CD2>Gal4, UAS-GFP). Immediately after heat shock, larvae were transferred at 18°C for 36-48 hours, then at 29°C for 36-48 hours to degrade Gal80. mCherry-Atg8a-positive structures were quantified using the PECAN image-analysis pipeline, separately within GFP-positive clones and in the surrounding GFP-negative wild-type tissue. The Atg8a-positive area was expressed as the percentage of tissue area covered by mCherry-Atg8a-positive structures. The ratio between Atg8a-positive coverage outside and inside the clones was calculated as a measure of the relative non-cell-autonomous autophagic response.

### 2.15 Statistical analysis

Statistical analyses were performed using GraphPad Prism 11. Data are presented as mean ± SEM or mean ± SD as specified for each experiment. Comparisons between two experimental groups were performed using unpaired two-tailed Student’s t-tests. Comparisons involving multiple genotypes or experimental conditions were performed using one-way ANOVA followed by Tukey’s multiple-comparisons test unless otherwise indicated. The number of biological replicates, animals, discs, or clones analyzed is indicated in the corresponding figure legends. Statistical significance was defined as P < 0.05. Significance is indicated as: ns, not significant; \**P* < 0.05; \*\**P* < 0.01; \*\*\**P* < 0.001; and \*\*\*\**P* < 0.0001.

## 3. Results

### 3.1 Myc promotes Gls-dependent accumulation of Atg8a-puncta in epithelial clones

Ammonia generated through glutaminolysis induces autophagy in mammalian cells. Given our previous findings that Myc increases Gls expression in *Drosophila* S2 cells and fat body [7, 8] and independently induces autophagy [18], we asked whether Gls-dependent glutamine catabolism contributes to Myc-induced autophagy by analyzing genetically defined clones in the wing imaginal disc, a system that allows both cell-autonomous and non-cell-autonomous autophagic responses to be examined within an intact epithelium. Quantitative RT-PCR analysis of RNA from wing imaginal discs revealed that Myc expression significantly increased *Gls* transcript levels (Figure 1A), along with changes in other genes involved in glutamine and amino acid metabolism (Supplementary Figure 2). To determine whether increased Gls functionally contributed to Myc-induced autophagy, we used clonal analysis to generate GFP-marked FLP-out clones expressing Myc alone or together with Gls-RNAi in animals carrying the mCherry-Atg8a reporter, expressed under the control of the endogenous Atg8a promoter [38]. Consistent with our previous analysis, Myc expression markedly increased the abundance of Atg8a-positive puncta compared with control clones (Figure 1C, E). In contrast, RNAi-mediated depletion of Gls significantly reduced the accumulation of Myc-induced Atg8a-positive puncta, restoring them to levels comparable to those of control clones (Figure 1C, F). Depletion of Gls alone had no effect on basal Atg8a puncta (Figure 1C, G). These findings demonstrate that Gls is required for the accumulation of Myc-induced Atg8a-puncta in the epithelial cells of the wing disc.

### 3.2 Ammonia promotes Atg8a-positive vesicle accumulation in Myc-expressing cells

To investigate whether ammonia contributes to the accumulation of the Atg8a puncta associated with Myc-dependent glutamine metabolism, we used *Drosophila* S2 cells carrying an HA-tagged Myc transgene under the control of a heat-shock-inducible promoter. Following heat shock, immunoblotting confirms robust Myc expression (Figure 2A). Consistent with our observations in wing imaginal discs, Myc significantly increased Gls transcription in S2 cells (Figure 2B). Because Gls-mediated glutaminolysis generates ammonia, we next measured ammonia released into the culture medium and found significantly higher levels in Myc-expressing cells (Figure 2C). We next tested whether this conditioned medium was sufficient to induce an autophagic response in *Drosophila* larval fat body, a well-established model for studying autophagy in which the large cell size facilitates visualization of Atg8a-positive structures. Fat bodies expressing the Atg8a-GFP reporter were dissected and incubated ex vivo with conditioned medium collected from control or Myc-expressing S2 cells. Conditioned medium from Myc-expressing cells induced a pronounced accumulation of Atg8a-GFP-positive vesicles compared with control medium (Figure 2D, E), indicating that Myc expression generated extracellular signals capable of stimulating an autophagic response.

Given the increased ammonia detected in the medium of Myc-expressing S2 cells, we next tested whether ammonia itself was sufficient to induce Atg8a-positive vesicles in the fat body. Ex vivo treatment with NH₄Cl induced Atg8a-positive puncta in a dose-dependent manner (Figure 2E), demonstrating that ammonia is sufficient to promote Atg8a-positive vesicle accumulation. We therefore examined the effect of exogenous NH₄Cl on Atg8a accumulation in clones in the wing imaginal disc and whether it could restore the reduced Atg8a phenotype caused by Gls depletion in Myc-expressing clones. NH₄Cl treatment alone induced the accumulation of mCherry-Atg8a-positive puncta in control wing discs (Figure 2J, compare the quantification in the first two lanes in grey), which was significantly increased when Myc was expressed (Figure 2G, J). Depletion of Gls significantly reduced the accumulation of Atg8a-positive puncta induced by Myc. NH₄Cl treatment significantly increased mCherry-Atg8a accumulation in Myc; Gls-RNAi clones, although not to the level observed with Myc alone, indicating a partial rescue of the phenotype (Figure 2I, J). Together, these results show that exogenous ammonia is sufficient to promote the accumulation of Atg8a-positive vesicles and partially overcome the effects of Gls depletion. The partial rescue supports a contribution of ammonia to the autophagic response associated with Myc-dependent glutamine metabolism, while suggesting that additional metabolic outputs associated with glutaminolysis may also contribute to this response.

### 3.3 MYC-induced autophagic flux is partially dependent on glutaminolysis

To determine whether the accumulation of Myc-induced Atg8a-positive puncta reflected productive autophagic flux rather than impaired turnover of autophagic structures, we employed the tandem mCherry-GFP-Atg8a reporter expressed in the wing pouch under the control of MS1096-Gal4 (Figure 3A, B). The tandem reporter distinguishes autophagosomes, which retain both GFP and mCherry fluorescence, from acidified autolysosomes, in which GFP fluorescence is quenched while mCherry remains stable. Control wing discs and discs expressing Myc alone or together with Gls-RNAi, or the corresponding RNAi, were analyzed (Figure 3D–G). Autophagic flux was quantified by measuring GFP/mCherry colocalization using Pearson’s correlation coefficient. In control wing discs, GFP and mCherry signals showed a high degree of colocalization (r ≈ 0.65). In contrast, Myc expression markedly reduced GFP/mCherry correlation (r ≈ 0.10), consistent with increased delivery of Atg8a-positive vesicles to acidic autolysosomes (Figure 3C, E). Concomitant depletion of Gls significantly increased the correlation coefficient to approximately 0.37, partially reversing the effect of Myc (Figure 3C, F). Gls depletion alone yielded a correlation coefficient of approximately 0.49, which was not significantly different from that of the corresponding control (Figure 3C, G). These results indicate that Myc promotes autophagic flux in the wing disc and that this response is partially dependent on Gls.

To independently assess autophagic activity, we analyzed endogenous Atg8a and Ref(2)P/SQSTM1 in S2 cells following induction of HA-Myc expression from a metallothionein promoter by addition of CuSO₄ to the culture medium for 4 h. Myc induction increased the abundance of lipidated Atg8a-II and the Atg8a-II/Atg8a-I ratio, while concomitantly reducing Ref(2)P protein levels (Figure 3H). Increased Atg8a lipidation together with reduced Ref(2)P levels is consistent with enhanced autophagic activity and cargo turnover in Myc-expressing S2 cells.

We next examined Ref(2)P turnover in vivo using a Ref(2)P-GFP reporter expressed under the control of the tubulin promoter. Consistent with the S2-cell results, Myc expression significantly reduced Ref(2)P-GFP fluorescence in the structures analyzed in the wing imaginal discs compared with controls (Figure 3I–J and K). Concomitant depletion of Gls restored Ref(2)P-GFP fluorescence to levels comparable to those of control discs, whereas Gls depletion alone did not significantly affect Ref(2)P-GFP levels (Figure 3K).

Together, these complementary approaches indicate that Myc-induced Atg8a puncta are associated with increased autophagic flux rather than impaired autophagic turnover. Moreover, the partial restoration of GFP/mCherry colocalization and recovery of Ref(2)P levels following Gls depletion support a contribution of Gls to Myc-induced autophagic flux.

### 3.4 Myc-induced autophagy requires Atg5 but is not suppressed by Rheb/TOR activation of atg1 depletion and Atg1

Previous studies have shown that ammonia generated during increased amino acid catabolism can induce an autophagic response that bypasses the canonical TOR-ULK1/Atg1 initiation pathway while retaining a requirement for the core autophagy machinery [44]. We therefore asked whether Myc-induced Atg8a accumulation shows a similar genetic activation. Thus, we examined the requirement of Rheb, Atg1, and Atg5 in Myc-induced autophagosome formation by monitoring mCherry-Atg8a-positive puncta in GFP-marked FLP-out clones (Figure 4). We first examined Rheb, an upstream activator of TOR signaling. Expression of the gain-of-function allele Rheb^AV4^ together with Myc did not suppress the Myc-induced increase in Atg8a puncta, and the levels were not significantly different from those observed with Myc alone (compare Figure 4B, E and Figure 4C, E). Rheb^AV4^ expression alone did not increase Atg8a puncta above control levels (Figure 4D, E). These results indicate that the accumulation of Atg8a-positive vesicles induced by Myc is not suppressed by activation of the Rheb/TOR axis. We next tested the requirement for Atg1/ULK1, the canonical autophagy-initiating kinase downstream of TOR. RNAi-mediated depletion of Atg1 did not suppress the increase in Atg8a-positive puncta induced by Myc, with Myc; Atg1-RNAi clones displaying levels comparable to those observed with Myc alone (Figure 4F–J). Atg1 depletion alone did not significantly alter the number of Atg8a-positive puncta compared with control clones. Thus, Atg1 depletion did not prevent Myc-induced Atg8a accumulation under these conditions.

**Figure 4.**
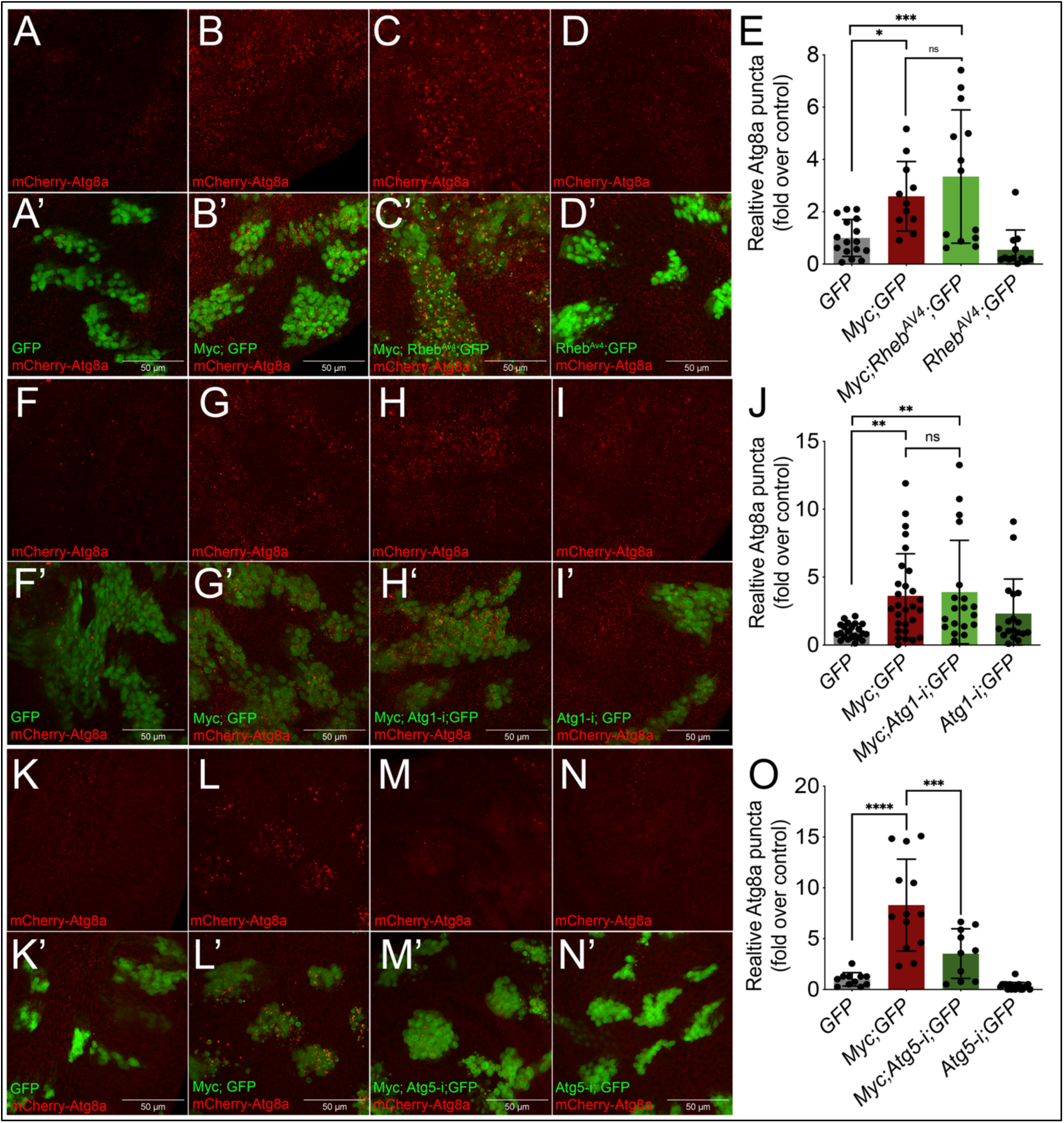
Myc-induced autophagy requires Atg5 but not Rheb or Atg1. (A–D′) Representative confocal images of GFP-marked FLP-out clones expressing the indicated transgenes in wing imaginal discs carrying the mCherry-Atg8a reporter: (A, A′) GFP control; (B, B′) Myc; GFP; (C, C′) Myc; Rheb^AV4^; GFP; and (D, D′) Rheb^AV4^; GFP. (E) Quantification of relative mCherry-Atg8a-positive puncta normalized to the GFP control. RhebAV4 did not suppress Myc-induced Atg8a accumulation. (F–I′) Representative images of (F, F′) GFP control; (G, G′) Myc; GFP; (H, H′) Myc; Atg1-i; GFP; and (I, I′) Atg1-i; GFP. (J) Quantification of relative mCherry-Atg8a-positive puncta. Atg1 depletion did not suppress Myc-induced Atg8a accumulation. (K–N′) Representative images of (K, K′) GFP control; (L, L′) Myc; GFP; (M, M′) Myc; Atg5-i; GFP; and (N, N′) Atg5-i; GFP. (O) Quantification of relative mCherry-Atg8a-positive puncta. Atg5 depletion significantly reduced Myc-induced Atg8a accumulation. In all images, mCherry-Atg8a is shown in red and GFP-marked clones in green. Each data point represents an individual FLP-out clone. Data were pooled from two or three independent biological experiments, with at least five to six wing discs analyzed per genotype. Bars represent mean ± SEM. Statistical significance was determined using one-way ANOVA followed by Tukey’s multiple-comparisons test. ns, not significant; *\*P* < 0.05; \*\**P* < 0.01; \*\*\**P* < 0.001; \*\*\*\**P* < 0.0001. Scale bars, 50 μm.

In contrast, depletion of the core autophagy component Atg5 significantly reduced the accumulation of Atg8a-positive puncta induced by Myc (Figure 4K-O). Myc; Atg5-RNAi clones displayed substantially fewer Atg8a-positive puncta than clones expressing Myc alone, whereas Atg5 depletion alone did not increase basal Atg8a accumulation.

These results indicate that Myc-induced autophagy requires Atg5 but is not suppressed by modulation of Rheb or Atg1, suggesting that it can proceed without strict dependence on the canonical Rheb–TOR–Atg1 pathway.

### 3.5 Myc is a critical determinant of Ras-driven tissue overgrowth

Metabolic interaction with the tumor microenvironment is an important feature of Ras-driven tumor growth [45]. We next examined the requirement for Myc in Ras^V12-^driven epithelial overgrowth using the *eyeless-FLP* system, which has previously been used to examine Myc-dependent growth [35]. This system generates genetically mosaic eye-antenna tissues expressing the transgenes of interest in either animals carrying *wild*-type Myc levels (*ey>dmyc^wt^*) or animals hemizygous for the hypomorphic *dmyc^P0^* allele (*ey>dmyc^P0^*), enabling quantitative analysis of growth phenotypes during larval development. Results from these experiments show that expression of *Ras^V12^; dlg-RNAi* (Figure 5A-B) or *Ras^V12^* (Figure 5F-G) in wild-type tissues induced massive neoplastic overgrowths associated with severe disruption of imaginal disc morphology (Figure 5B-G and graphs E and J) and larval lethality. Strikingly, reducing Myc activity with the *dmyc^P0^* allele markedly suppressed tissue overgrowth in both models (Figure 5D and I, and graphs E and J), restoring disc size to levels close to those of control tissues (Figure 5C and H, and graphs E and J, respectively). Together, these findings demonstrate that Myc is required for Ras-driven epithelial overgrowth and that this requirement is also observed with Ras^V12^ alone, independently of *dlg* depletion.

**Figure 5.**
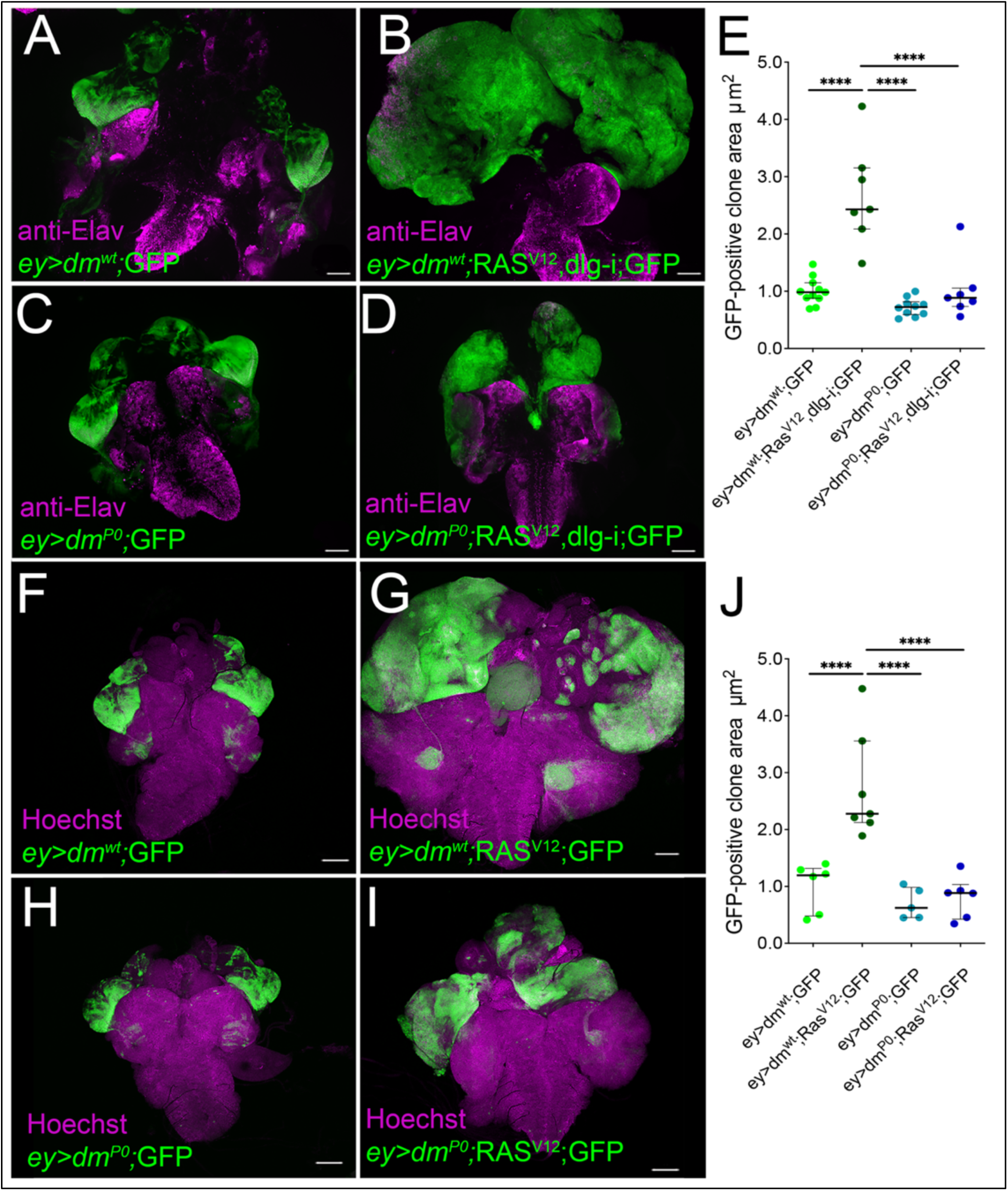
Myc is required for Ras^V12-^induced overgrowth in eye-antenna imaginal discs. (A–D) Representative confocal images of third-instar larval eye-antennal imaginal discs with GFP-marked clones generated within the *eyeless* domain using the *ey-FLP* system. GFP-positive tissues are shown in green, and Elav staining, marking differentiated photoreceptor cells, is shown in magenta. (A) *ey>dm ^WT^*; GFP control discs. (B) *ey>dm ^WT^; Ras^V12^, dlg-RNAi; GFP* discs exhibiting marked tissue overgrowth. (C) *ey>dm^P0^; GFP* control discs. (D) *ey>dm^P0^; Ras^V12^, dlg-RNAi; GFP* discs, in which tumor overgrowth is strongly suppressed by reduced Myc activity. (E) Quantification of the GFP-positive clone area for the genotypes shown in A–D. (F–I) Representative confocal images from an independent experiment examining Ras^V12^-driven overgrowth in the absence or presence of reduced Myc activity. GFP-positive clones are shown in green, and nuclei stained with Hoechst 33352 are shown in magenta. (F) *ey>dm ^WT^*; GFP, (G) *ey>dm^WT^; Ras^V12^; GFP* (H) *ey>dm^P0^; GFP*, and (I) *ey>dm^P0^; Ras^V12^; GFP*. (J) Quantification of the GFP-positive clone area for the genotypes shown in F-I. Data are normalized to the mean area of control *ey>dm ^WT^*; GFP discs. Each point represents an individual animal; bars indicate mean ± SD. Statistical significance was determined by one-way ANOVA with multiple-comparison correction. *\*\*\*P < 0.001; ****P < 0.0001*. Scale bars, 100 μm.

### 3.6 Myc-dependent glutamine metabolism in Ras^V12^-driven tumor models promotes non-cell-autonomous autophagy

We next asked whether the Myc-Gls-dependent autophagic response identified above extends to Ras-transformed epithelial cells. Because Ras^V12^ clones frequently displayed substantial variability in growth, potentially reflecting competitive interactions with surrounding wild-type tissue [25, 46], we used a Gal80^ts^ to temporally control Ras^V12^ expression, thereby generating more uniform and reproducible clones. To assess the contribution of Myc and Gls to autophagy during Ras^V12^-driven clonal overgrowth, without the additional effect of *dlg* depletion, we first measured Myc protein levels. Myc levels were comparable in Ras^V12^ and Ras^V12^; dlg-RNAi clones (Supplementary Figure 3), indicating that *dlg* depletion did not alter Myc expression. We therefore focused our analysis on Ras^V12^ alone. Temporally controlled clonal analysis revealed that Atg8a-positive puncta remained low within Ras^V12^ clones and were comparable to levels observed in control clones (Figure 6; compare A with C and G with I). In contrast, a marked accumulation of Atg8a-positive vesicles was observed in the surrounding wild-type tissue (Figures 6C and I), revealing a non-cell-autonomous autophagic response. Consistent with this observation, mCherry-Atg8a coverage was increased outside Ras^V12^ clones (Figures 6E and K), resulting in a higher ratio of Atg8a coverage outside versus inside the clones (Figures 6F and L). A similar non-cell-autonomous pattern was observed in Ras^V12^; dlg-RNAi clones (Supplementary Figure 4).

**Figure 6.**
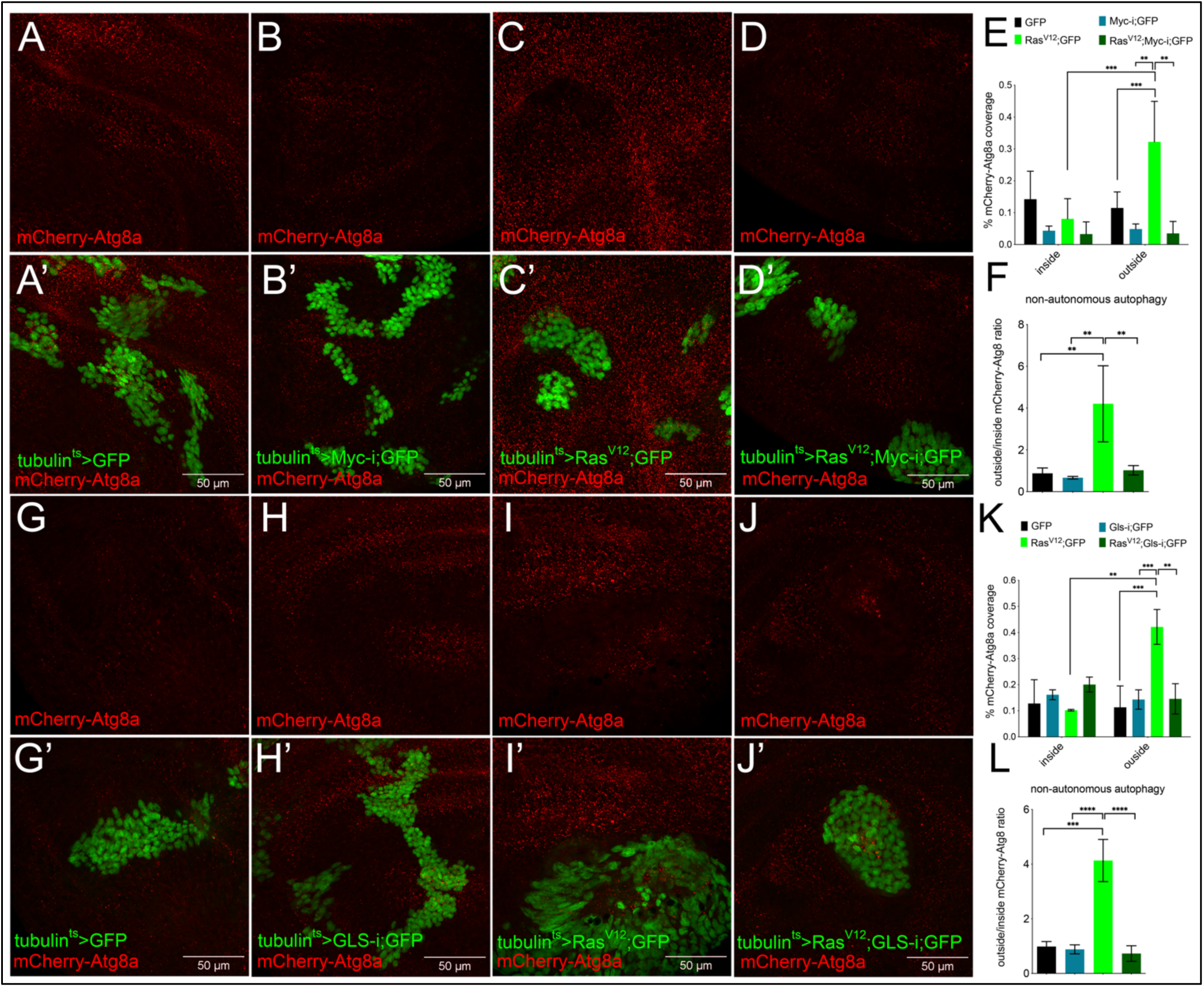
Myc and Gls are required for non-cell-autonomous autophagy surrounding Ras^V12^ clones. (A–D′) Representative confocal images of wing imaginal discs carrying the mCherry-Atg8a reporter and GFP-marked FLP-out clones of the indicated genotypes: (A, A′) control; (B, B′) Myc-RNAi; (C, C′) Ras^V12^; and (D, D′) Ras^V12^; Myc-RNAi. mCherry-Atg8a is shown in red and GFP-marked clones in green. (E) Quantification of mCherry-Atg8a-positive area inside and outside clones, expressed as a percentage of area coverage. (F) Ratio of mCherry-Atg8a coverage outside versus inside clones. Ras^V12^ increased Atg8a accumulation in the surrounding tissue relative to that within the clones, an effect reduced by Myc depletion. (G–J′) Representative images of (G, G′) control; (H, H′) Gls-RNAi; (I, I′) Ras^V12^; and (J, J′) Ras^V12^; Gls-RNAi clones. (K) Quantification of mCherry-Atg8a-positive area inside and outside clones. (L) Ratio of mCherry-Atg8a coverage outside versus inside clones. Gls depletion reduced the non-cell-autonomous accumulation of Atg8a associated with Ras^V12^ clones. Statistical significance was determined using one-way ANOVA followed by Tukey’s multiple-comparisons test. Error bars represent SD. Scale bars, 50 μm.

To determine whether Myc contributes to this non-cell-autonomous response, we reduced Myc levels in Ras^V12^ clones. Myc depletion markedly reduced the accumulation of Atg8a-positive vesicles in the surrounding tissue (Figures 6D–F), indicating that Myc activity within Ras^V12^ cells contributes to autophagy in neighboring wild-type cells. We next asked whether Gls, which mediates Myc-dependent glutamine metabolism, participates in this response. Gls depletion similarly reduced Atg8a-positive vesicle accumulation in the tissue surrounding Ras^V12^ clones (Figures 6J–L), supporting a role for Myc-dependent glutamine metabolism in promoting non-cell-autonomous autophagy.

Together, these findings identify Myc-dependent glutamine metabolism as a determinant of the non-cell-autonomous autophagic response elicited by Ras^V12^-transformed cells, linking metabolic activity within transformed cells to autophagy in the surrounding tissue.

## 4. Discussion

MYC-driven growth requires coordinated changes in cellular metabolism and stress-adaptive pathways that support cellular fitness. Here, we identify glutamine metabolism as an important component of the autophagic response to Myc, linking Gls-dependent glutaminolysis to productive autophagic flux. This relationship extends beyond the cell-autonomous effects of Myc. In Ras^V12^-transformed epithelia, depletion of either Myc or Gls within transformed cells markedly reduced autophagy in neighboring wild-type tissue, linking the metabolic state of Ras-transformed cells to a non-cell-autonomous autophagic response.

MYC-dependent glutamine utilization is generally considered to sustain the bioenergetic and biosynthetic demands of proliferating cells [3, 6]. Our findings extend this function by linking MYC-dependent glutamine metabolism to the regulation of autophagy. Myc affected multiple components of glutamine and amino acid metabolism (Supplementary Figure 1), with Gls emerging as a functionally relevant mediator of this response. Gls depletion suppressed Myc-induced Atg8a accumulation and impaired autophagic flux, supporting a requirement for glutaminolysis to sustain productive autophagy rather than simply promoting the accumulation of autophagic structures (Figures 1 and 3). This interpretation is further supported by changes in Atg8a lipidation and Ref(2)P/SQSTM1 turnover. Notably, the effect of Myc on Ref(2)P was tissue dependent. Myc reduced Ref(2)P levels in wing imaginal discs and S2 cells, consistent with increased cargo turnover, whereas Ref(2)P accumulated in the larval fat body (Supplementary Figure 5), in agreement with previous observations in this tissue [17]. These differences suggest that the relationship between Myc activity and Ref(2)P turnover is tissue dependent, potentially reflecting the distinct metabolic and autophagic functions of the tissues examined. Our findings extend previous studies that link elevated Myc activity to autophagy via distinct cellular stress pathways. In *Drosophila,* Myc-induced proteotoxic stress activates the UPR and PERK-dependent autophagy together with the Nrf2 antioxidant response [17], whereas alterations in lipid metabolism contribute to Myc-dependent autophagy and tissue growth [18]. A link between MYC-dependent glutamine metabolism and autophagy has also been reported in prostate cancer, where MYC-driven GLS activity promotes ATG5-dependent autophagy, contributing to oxidative stress control, stem-like properties, and treatment resistance [14].

Our results identify glutamine metabolism as an additional component of the autophagic response to MYC and suggest that distinct metabolic and proteostatic pathways may converge on the autophagic machinery to meet the demands imposed by elevated MYC activity.

A potential mechanistic link between glutaminolysis and autophagy is provided by ammonia, a by-product of glutamine catabolism that can act as a diffusible inducer of autophagy in mammalian cells [9, 10]. Consistent with this model, Myc increased ammonia production in S2 cells, whereas exogenous NH₄Cl promoted accumulation of Atg8a-positive vesicles and partially restored this response following Gls depletion (Figure 2). The partial rescue by NH₄Cl supports ammonia as one metabolic output of Myc-dependent glutaminolysis that contributes to the autophagic response, while also indicating that additional consequences of glutamine catabolism are likely to participate.

Interestingly, the genetic requirements of Myc-induced autophagy show similarities to those described for ammonia-induced autophagy in mammalian cells. In wing imaginal discs, depletion of Atg1 did not suppress Myc-induced Atg8a accumulation, whereas depletion of Atg5 strongly reduced this response (Figure 4), indicating that Myc-induced autophagy requires Atg5 but not Atg1. Moreover, activation of Rheb/TOR signaling failed to suppress Myc-induced Atg8a accumulation. Ammonia-induced autophagy in mammalian cells can similarly proceed independently of ULK1/ULK2 while retaining a requirement for ATG5 [44]. These parallels support the possibility that Myc-dependent glutaminolysis engages an autophagic program that can operate independently of canonical TOR-Atg1 regulation while retaining a requirement for the core autophagy machinery. This does not exclude context-dependent contributions of Atg1 or other autophagy regulators, as previous studies have implicated Atg1 and additional autophagy components in Myc-dependent growth and stress responses in *Drosophila* [17].

We next extended these findings to Ras-transformed epithelia. Reducing Myc markedly suppressed Ras^V12^-driven epithelial overgrowth, both alone and in combination with dlg-RNAi (Figure 5), supporting a requirement for Myc in Ras-driven tissue expansion. Analysis of autophagy in temporally controlled Ras^V12^ clones revealed a striking spatial response. Atg8a-positive puncta remained low within Ras^V12^ clones and were comparable to those observed in control clones, whereas Atg8a-positive structures accumulated prominently in the surrounding wild-type epithelium (Figure 6). Previous work in *Drosophila* has shown that Ras^V12^; scrib-/-tumors induce non-cell-autonomous autophagy in neighboring and distant tissues, which supports tumor growth by increasing nutrient availability to transformed cells [33, 34].

Our findings extend this concept by addressing the opposite side of this metabolic interaction: how the metabolic state of transformed cells contributes to the induction of autophagy in the surrounding tissue. Depletion of either Myc or Gls within Ras^V12^ clones markedly reduced Atg8a accumulation in neighboring wild-type cells, indicating that Myc-dependent glutamine metabolism in transformed cells contributes to this non-cell-autonomous autophagic response. How metabolic information is transmitted from Ras^V12^ cells to the surrounding epithelium remains to be established. Ammonia is an attractive candidate because of its diffusible nature and because both conditioned medium from Myc-expressing cells and exogenous NH₄Cl induced Atg8a-positive structures in our experimental systems. However, our experiments do not establish that ammonia is the signal responsible for autophagy surrounding Ras^V12^ cells, and other Gls-dependent metabolic outputs may contribute. It will also be interesting to determine whether autophagy in neighboring cells engages the canonical TOR-dependent program associated with nutrient deprivation or instead reflects a response to metabolic signals generated by Ras^V12^-transformed cells.

Together, our findings identify Gls-dependent glutaminolysis as a component of Myc-induced autophagy and link this metabolic program to non-cell-autonomous autophagy surrounding Ras-transformed cells. The Atg5 dependence, lack of requirement for Atg1, and resistance to Rheb/TOR activation support an autophagic response distinct from canonical TOR-Atg1 regulation, with ammonia representing one potential metabolic contributor. These findings extend the role of MYC-dependent glutamine metabolism beyond its established anabolic functions and reveal a connection between the metabolic state of transformed cells and autophagy in the surrounding tissue.

## 5. Conclusion

In conclusion, our findings identify glutamine metabolism as a link between MYC activity and autophagic signaling in epithelial tissues. Gls-dependent glutaminolysis not only contributes to the cell-autonomous response to MYC but also influences autophagy in cells surrounding Ras-transformed epithelia. These findings broaden the role of MYC-dependent metabolic signaling beyond the transformed cell and suggest that glutaminolysis-driven metabolic communication can help shape the autophagic response within the local tumor microenvironment.

## Appendix A

Supplementary information, Figures 1-5 and a Table 1

## CRediT authorship contribution statement

**FD:** Writing, Investigation, Formal analysis. **LC:** Investigation, Formal analysis, Writing-review and editing. **SB:** Investigation, Writing-review and editing. **VM:** Methodology, Investigation. **PB:** Conceptualization, Supervision, Funding acquisition, Project administration, Investigation, Writing original draft, Writing-review and editing.

## Funding acquisition

This work was supported by grants from the NIH-NIDDK-DK085047 for P.B, and by an institutional fellowship to FD and VM.

## Acknowledgments

We thank Juhasz Gabor, Tor Erik Rusten, Thomas Neufeld, and Hugo Stocker for stock lines, the Bloomington Stock Center (NIH P40OD018537). We are grateful that the Department CIBIO Core Facilities is supported by the European Regional Development Fund (FESR) 2021–2027. The Dipartimento di Eccellenza 2023-2027, Legge 232/2016, project n 40613, funded by the MUR. This document includes language and clarity improvements made using ChatGPT (OpenAI).

