## Supplementary and table 1 for "Glutaminase contributes to MYC-induced cell-autonomous autophagy and to Ras^V12^-dependent non-autonomous autophagy in the Drosophila wing disc epithelium"

Supplementary Figure 1

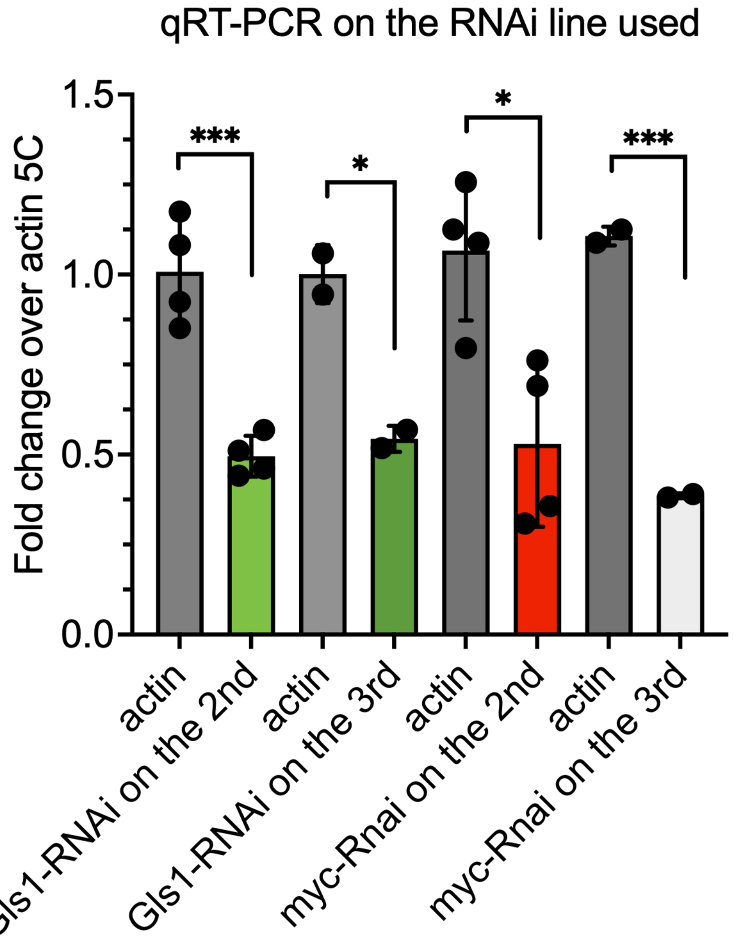


**Figure 1. Validation of RNAi lines used in this study by qRT-PCR.** Knockdown efficiency of the independent Gls1-RNAi and myc-RNAi lines used in this study was assessed by qRT–PCR. Transcript levels were normalized to Actin5C and are expressed as fold change relative to the corresponding control. Bars represent mean ± SD; individual biological replicates are shown as dots. Statistical significance: *P < 0.05; ***P < 0.001.

Supplementary Figure 2

Myc-induced gene expression:

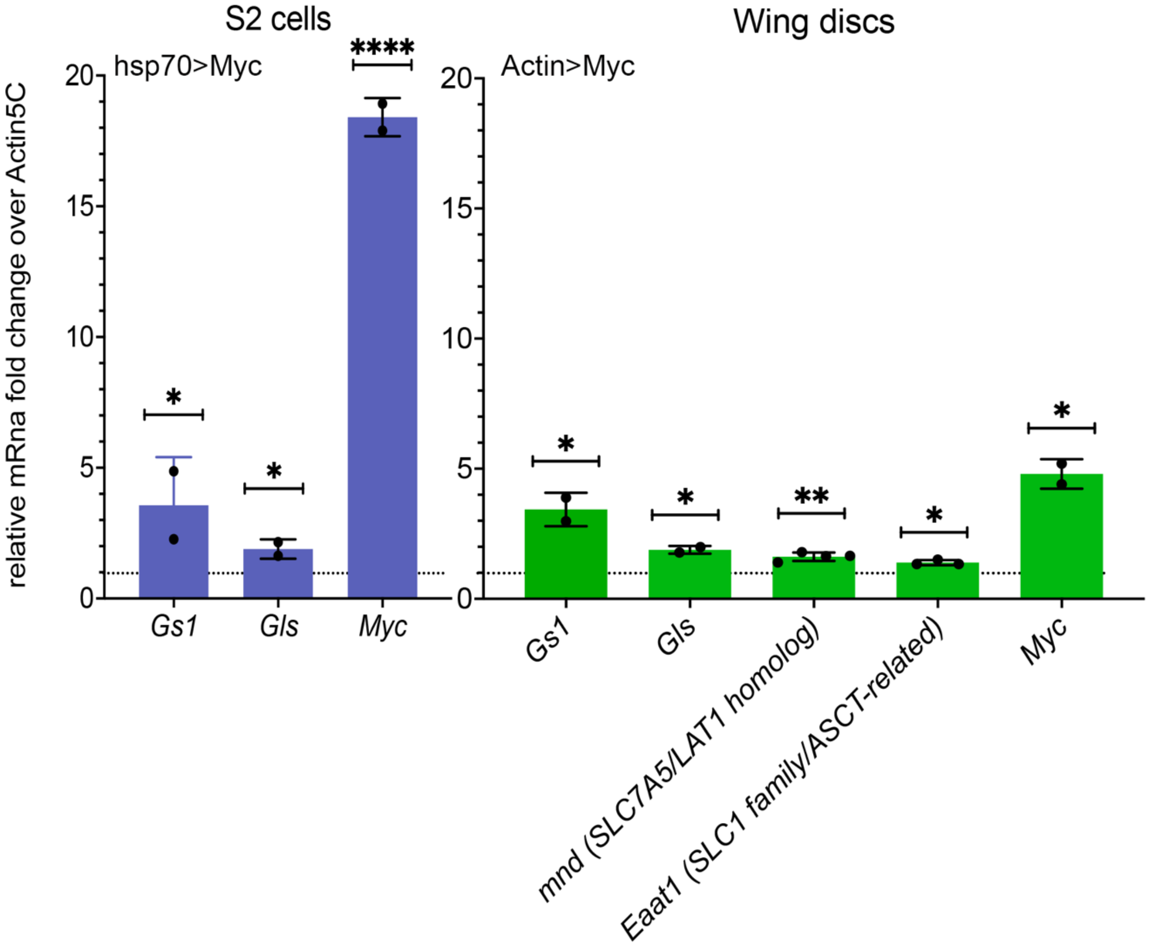


**Figure 2. Myc induces the expression of genes involved in glutamine and amino acid metabolism.** qRT-PCR analysis of the indicated genes in Myc-overexpressing S2 cells and wing imaginal discs. In S2 cells, Myc expression was induced from the hsp70 -Gal4 promoter by heat shock for 30 min at 37 °C, followed by 2 h of recovery before RNA extraction. In wing imaginal discs, Myc was ubiquitously expressed using the Actin-Gal4 promoter, and discs were collected from third-instar larvae. RNA was extracted and processed as described in Materials and Methods. Transcript levels were normalized to Actin5C and expressed as fold change relative to the corresponding control using the 2^-ΔΔCt method, with the control set to 1. Data are shown as mean ± SEM of ≥3 independent biological replicates. Statistical significance was determined using Student's t-test. *P < 0.05; **P < 0.01; ****P < 0.0001. S2 cells and wing imaginal discs represent independent experimental datasets and were analyzed separately.

Supplementary Figure 3

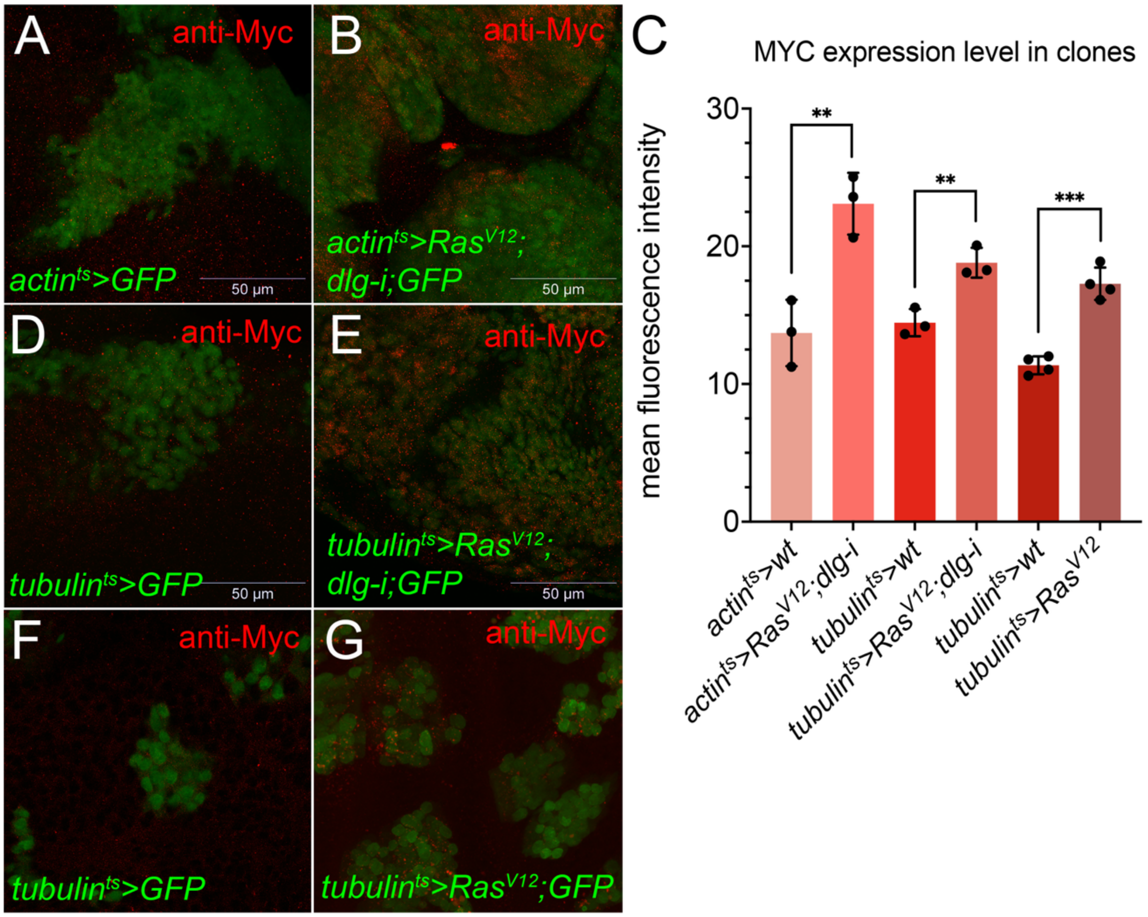


**Figure 3. Myc protein levels increase in $Ras^{V12}$ and $Ras^{V12}$; dlg-RNAi clones.**

(A, B) Representative confocal images of GFP-marked FLP-out clones generated using the actin promoter: (A) control and (B) $Ras^{V12}$; dlg-RNAi. (D–G) Representative images of GFP-marked FLP-out clones generated using the tubulin promoter: (D) control, (E) $Ras^{V12}$; dlg-RNAi, (F) control, and (G) $Ras^{V12}$. Myc protein was detected by immunofluorescence (red), and GFP-marked clones are shown in green. (C) Quantification of mean Myc fluorescence intensity within GFP-positive clones. Myc levels were significantly increased in both $Ras^{V12}$ and $Ras^{V12}$; dlg-RNAi clones relative to their corresponding controls. Data are shown as mean ± SEM; each point represents an independent biological replicate. Statistical significance was determined using an unpaired two-tailed Student's t-test; $**P < 0.01$; $***P < 0.001$. Scale bars, 50 µm.

Supplementary Figure 4

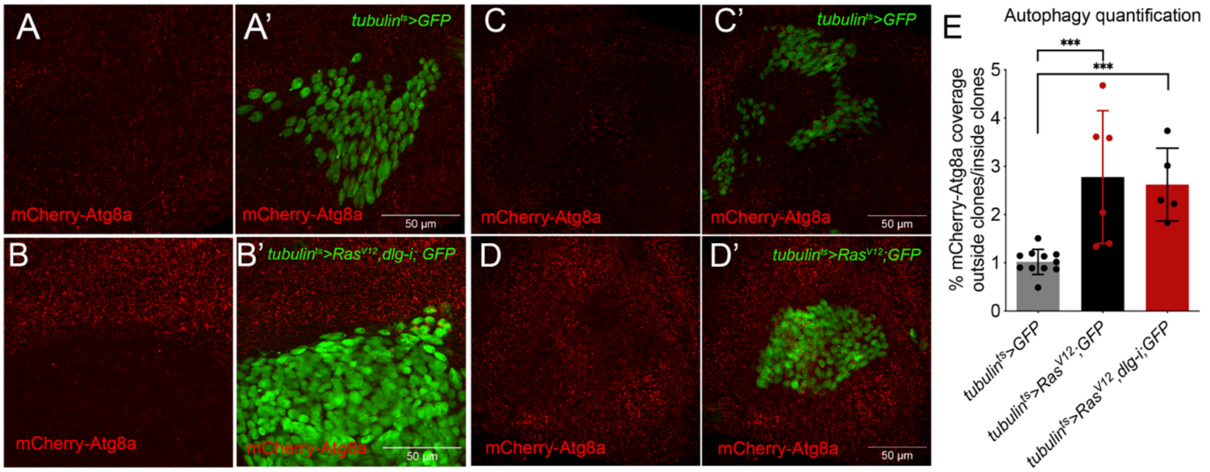


**Figure 4. Ras$^{V12}$ and Ras$^{V12}$; dlg-RNAi clones induce comparable non-cell-autonomous Atg8a accumulation.** (A–D′) Representative confocal images of wing imaginal discs carrying the mCherry-Atg8a reporter and GFP-marked FLP-out clones generated using the tubulin promoter: (A, A′) control GFP, (B, B′) Ras$^{V12}$; dlg-RNAi; GFP, (C, C′) control GFP, and (D, D′) Ras$^{V12}$; GFP. mCherry-Atg8a is shown in red and GFP-marked clones in green. (E) Quantification of mCherry-Atg8a coverage outside relative to inside the clones. Both Ras$^{V12}$ and Ras$^{V12}$; dlg-RNAi clones showed significantly increased non-cell-autonomous Atg8a accumulation compared with control clones, with comparable responses between the two Ras$^{V12}$ models. Data are shown as mean ± SD; each point represents an individual clone. Statistical significance was determined using one-way ANOVA followed by Tukey's multiple-comparisons test; ***P < 0.001. Scale bars, 50 µm

Supplementary Figure 5

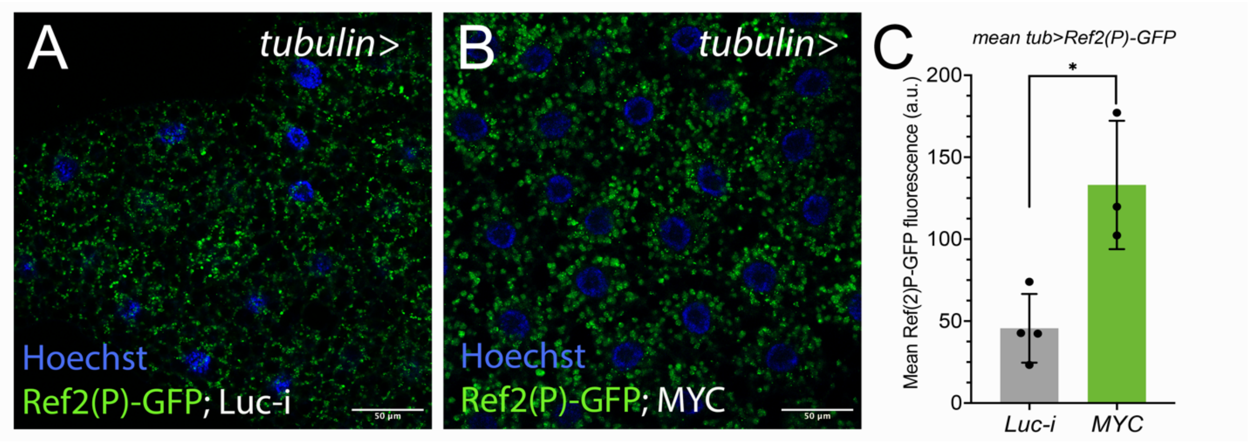


**Figure 5: Myc induces Ref(2)P-GFP accumulation in larval fat body.** (A, B) Representative confocal images of larval fat bodies expressing the constitutive *tubulin>Ref(2)P-GFP* reporter and control UAS-Luc-I (A) or UAS-HA-Myc (B). Nuclei were stained with Hoechst (blue). Scale bars, 50 µm. (C) Quantification of background-corrected mean Ref(2)P-GFP fluorescence

intensity in fat bodies of animals of the indicated genotype. Multiple regions of interest (ROIs) were measured for each fat body and averaged to obtain one value per biological sample. Bars represent the mean ± SD, and each dot corresponds to one independent fat body. Statistical significance was assessed using an unpaired two-tailed Student's t-test; $P < 0.05$ (*). The observed increase in Ref(2)P-GFP fluorescence is consistent with previous reports of increased endogenous Ref(2)P protein in MYC-expressing fat bodies [1]

[1] P. Nagy, A. Varga, K. Pircs, K. Hegedus, G. Juhasz, Myc-driven overgrowth requires unfolded protein response-mediated induction of autophagy and antioxidant responses in Drosophila melanogaster, PLoS Genet 9(8) (2013) e1003664.
[2] F. Parisi, S. Riccardo, S. Zola, C. Lora, D. Grifoni, L.M. Brown, P. Bellosta, dMyc expression in the fat body affects DILP2 release and increases the expression of the fat desaturase Desat1 resulting in organismal growth, Dev Biol 379(1) (2013) 64–75.

**Table 1: Sequences of the primers used in this work.**

| Target | Sequence | Reference |
| --- | --- | --- |
| *Gls* | F: TCGTCGTCATTAGGGCAAC | [2] |
| | R: CGCTTCAAAGATCCGTCTC | |
| *Gs1* | F: TGCGTCTGCTGCGTACTGGC | [2] |
| | R: CGGCGTTTCCAGGTTGCGGTA | |
| *Myc* | F: CATAACGTCGACTTGCGTG | [2] |
| | R: GAAGCTCCCTGCTGATTTGC | |
| *Mnd/SLC7A/LAT1 homolog* | F: AGTCAGAGCCGTAATCGCAT | This work |
| | R: ATAGGCACTGCATTTGGTCC | |
| *Eaat1/SLC/ASCT related* | F: GGACAACATGGGCATCGATC | This work |
| | R: GAT TCCAGCAGCTCCAATCG | |
